# Tetrad analysis without tetrad dissection: Meiotic recombination and genomic diversity in the yeast *Komagataella phaffii (Pichia pastoris)*

**DOI:** 10.1101/704627

**Authors:** Stephanie Braun-Galleani, Julie A. Dias, Aisling Y. Coughlan, Adam P. Ryan, Kevin P. Byrne, Kenneth H. Wolfe

**Affiliations:** UCD Conway Institute, School of Medicine, University College Dublin, Dublin, Ireland; Department of Mathematics and Statistics, McGill University, Montreal, Quebec, Canada

## Abstract

*Komagataella phaffii* is a yeast widely used in the pharmaceutical and biotechnology industries, and is one of the two species that were formerly called *Pichia pastoris*. However, almost all laboratory work on *K. phaffii* has been done on strains derived from a single natural isolate, CBS7435. There is little information about the genetic properties of *K. phaffii* or its sequence diversity. Genetic analysis is difficult because, although *K. phaffii* makes asci with four spores, the spores are small and tend to clump together, making the asci hard to dissect. Here, we sequenced the genomes of all the known isolates of this species, and find that *K. phaffii* has only been isolated from nature four times. We analyzed the meiotic recombination landscape in a cross between auxotrophically marked strains derived from two isolates that differ at 44,000 single nucleotide polymorphism sites. We conducted tetrad analysis by making use of the property that haploids of this species do not mate in rich media, which enabled us to isolate and sequence the four types of haploid cell that are present in the colony that forms when a tetratype ascus germinates. We found that approximately 25 crossovers occur per meiosis, which is 3.5 times fewer than in *Saccharomyces cerevisiae*. Recombination is suppressed, and genetic diversity among natural isolates is low, in a region around centromeres that is much larger than the centromeres themselves. Our method of tetrad analysis without tetrad dissection will be applicable to other species whose spores do not mate spontaneously after germination.

**Author summary:** To better understand the basic genetics of the budding yeast *Komagataella phaffii*, which has many applications in biotechnology, we investigated its genetic diversity and its meiotic recombination landscape. We made a genetic cross between strains derived from two natural isolates, and developed a method for characterizing the genomes of the four spores resulting from meiosis, which were previously impossible to isolate. We found that *K. phaffii* has a lower recombination rate than *Saccharomyces cerevisiae*. It shows a large zone of suppressed recombination around its centromeres, which may be due to the structural differences between centromeres in *K. phaffii* and *S. cerevisiae*.

## Introduction

*Komagataella phaffii* is the most widely used yeast species for production of heterologous proteins, such as the expression of antibody fragments for the pharmaceutical industry. It has several advantages over *Saccharomyces cerevisiae* as a cell factory, including thermotolerance, respiratory growth to very high cell densities, and the ability to induce high levels of foreign gene expression using its methanol oxidase promoter [1–3]. *K. phaffii* is better known under its previous name *Pichia pastoris*, but in 2009 it was realized that the name ‘*P. pastoris*’ had been applied to a heterogeneous group of strains that actually belong to two separate species, which are now called *K. phaffii* and *K. pastoris* [4]. Their genomes differ by approximately 10% DNA sequence divergence and two reciprocal translocations [5]. Phylogenetically, *Komagataella* species are members of the methylotrophic yeasts clade (family Pichiaceae) and are only distantly related to better-known yeasts such as *S. cerevisiae* and *Candida albicans* [6].

Almost all research on *K. phaffii* has been done using the genetic background of strain CBS7435 (synonymous with NRRL Y-11430) [7]. The origin of this strain has been unclear because it was deposited in the CBS and NRRL culture collections in connection with a US patent granted to Phillips Petroleum (see discussion in [4]), but in this study we show that CBS7435 is identical to the type strain of *K. phaffii*, which was isolated from an oak tree. The widely-used *K. phaffii* strains GS115 and X-33 are derivatives of CBS7435 and are components of a commercial protein expression kit marketed by Invitrogen / Life Technologies / Thermo Fisher. GS115 was made from CBS7435 by random mutagenesis with nitrosoguanidine and includes a *his4* mutation among a few dozen point mutations [5, 8]. X-33 is a derivative of GS115 in which the *HIS4* gene was reverted to wildtype by site-directed mutagenesis [9].

The public culture collections include a few natural isolates of *K. phaffii*, but these have received little attention. Most of them were isolated from exudates (slime fluxes) on trees. The genetic diversity that is naturally present in populations of *K. phaffii* could potentially be used to improve its performance in biotechnological applications. In principle, beneficial alleles from natural isolates could be introduced into biotechnology strains by breeding. As a first step towards this goal, in this study we surveyed the nucleotide sequence diversity that is present in all six isolates of *K. phaffii* that are available from culture collections.

Although techniques for inserting foreign genes into the *K. phaffii* genome and controlling their expression are well established, other aspects of the genetics and life cycle of this yeast are much less studied [10]. *K. phaffii* has four chromosomes and grows primarily as a haploid. Mating only occurs when induced by nitrogen depletion [10–12]. Zygotes usually sporulate immediately after mating, but in crosses between haploids carrying auxotrophic markers, the diploid progeny can be maintained by transferring them to nitrogen-replete media and selecting for prototrophy [11]. Mating occurs between *MAT***a** and *MAT*α cells, and haploids can switch their mating type by inverting a 138-kb section of chromosome 4 [12, 13]. The recent development of stable heterothallic strains of *K. phaffii* makes it possible to carry out controlled genetic crosses without requiring selectable markers [13, 14].

One of the classical techniques used for genetic studies in yeasts, particularly for analysis of recombination and gene conversion, is tetrad analysis [15–18]. Yeast meiosis usually results in the formation of an ascus containing four ascospores (a tetrad), which carry the four sets of chromosomes produced by meiosis. In species such as *S. cerevisiae* or *Schizosaccharomyces pombe*, the ascus can be dissected by micromanipulation, allowing each spore to be germinated separately into a haploid colony (a segregant) and analyzed. Tetrad analysis is also possible in other eukaryotes in which the four products of a meiosis remain attached to each other, such as other fungi, green algae, and plants, although recovery of the four meiotic products in plants for whole genome sequencing is difficult [19–22].

In the yeasts *S. cerevisiae*, *Saccharomyces paradoxus*, and *Lachancea kluyveri*, the meiotic recombination landscape has been studied in detail by analyzing the genomes of the four segregants from multiple tetrads, either by genome sequencing or by microarray analysis of dissected tetrads [23–27]. However, tetrad analysis has not been achieved in *K. phaffii* because its spores are much smaller than those of *S. cerevisiae* and they tend to stick together and form clumps [10]. *K. phaffii* tetrads are 1-2 μm in diameter (**Fig 1**), which is approximately four times smaller than tetrads in *S. cerevisiae* [28, 29]. Previous genetic analysis in *K. phaffii* has relied on random spore methods, which are less powerful than tetrad analysis [10]. Similar problems make tetrad dissection in the related methylotrophic yeast *Ogataea polymorpha* difficult but not impossible [30, 31]. Here, we developed a method for tetrad analysis in *K. phaffii* that does not require asci to be dissected. We made a genetic cross between strains derived from CBS7435 and a divergent natural isolate, enabling us to investigate the landscape of meiotic recombination (that is, the distribution of recombination sites along chromosomes) in this species for the first time.

**Fig 1.**
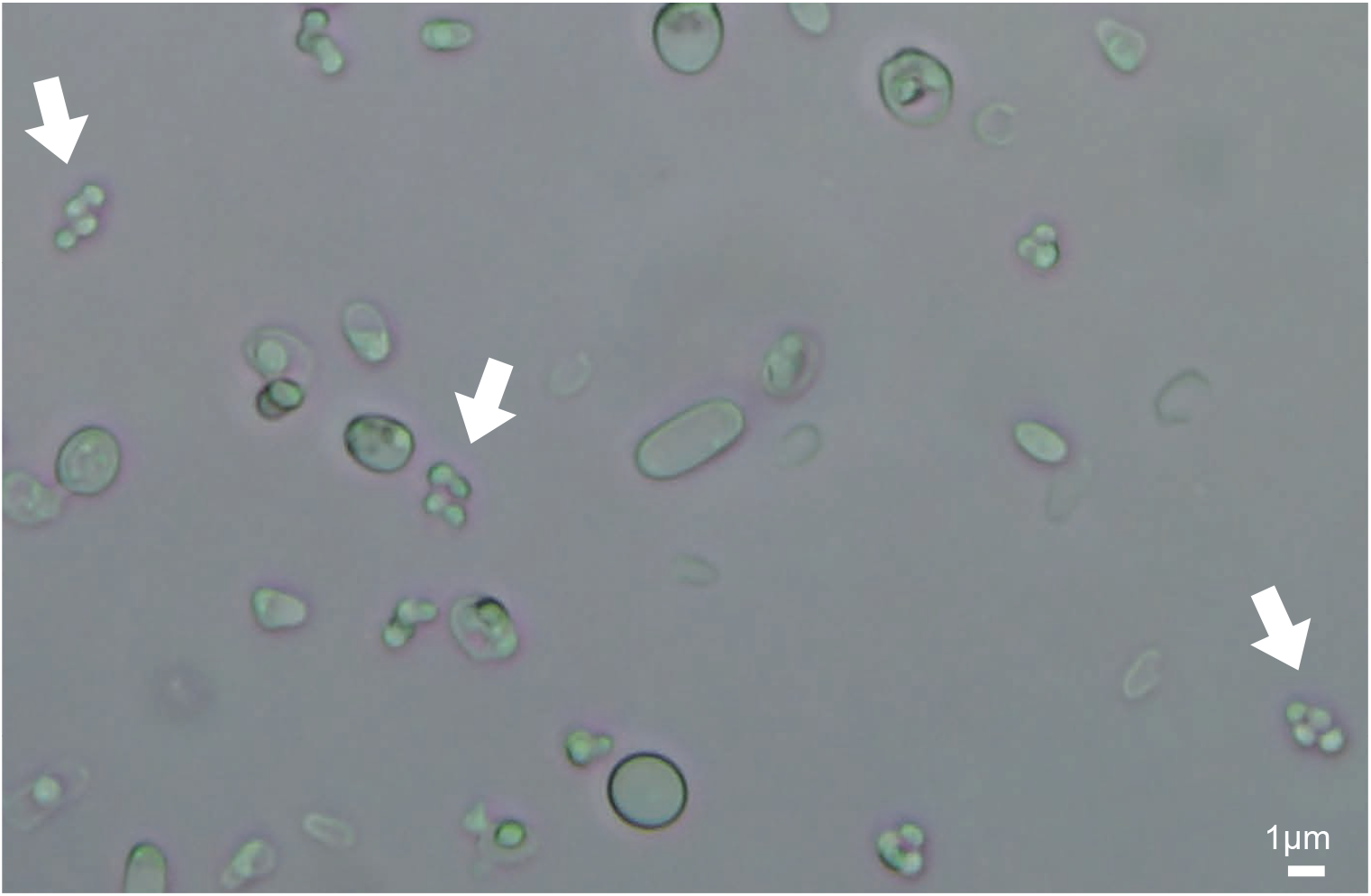
Bright field microscope image of a sporulated *K. phaffii* culture. Asci containing four spores are indicated with white arrows. Scale bar, 1 μm. This image is from a cross of CBS7435 x Pp2.

## Results

### Nucleotide diversity in natural isolates of *K. phaffii*

We obtained five isolates of *K. phaffii* from the NRRL culture collection and sequenced their genomes. For convenience we refer to them as Pp1 – Pp5; their strain numbers in the NRRL and CBS culture collections are given in **Table 1**. As far as we know, no other isolates of this species are available from public culture collections apart from CBS7435 [4, 32]. We mapped the Illumina reads from each strain to the CBS7435 reference genome sequence [33] using BWA, and identified variant sites using the GATK SNP (single nucleotide polymorphism) calling pipeline. We included strain GS115 in this analysis, using the genome sequence reported by Love *et al*. [5]. A total of 64,019 variable SNP sites were identified among the six strains relative to CBS7435 (Table 1), and a phylogenetic tree of the strains was constructed from the variable sites (**Fig 2**).

**Table 1.**
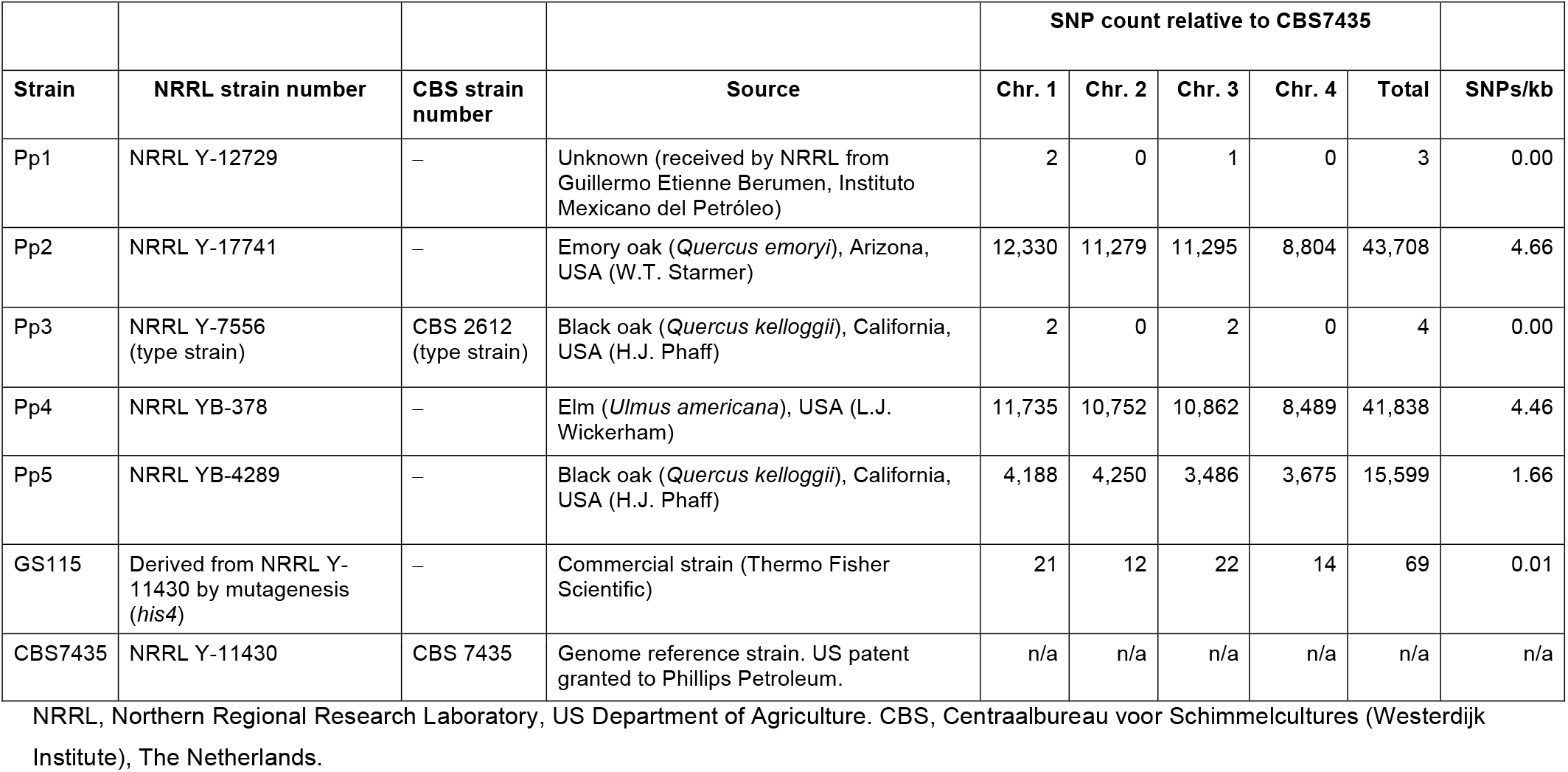
*Komagataella phaffii* strains used in this study, and numbers of SNPs identified on each chromosome relative to CBS7435.

| Strain | NRRL strain number | CBS strain number | Source | SNP count relative to CBS7435 |  |  |  |  | SNPs/kb |
| --- | --- | --- | --- | --- | --- | --- | --- | --- | --- |
|  |  |  |  | Chr. 1 | Chr. 2 | Chr. 3 | Chr. 4 | Total |  |
| Pp1 | NRRL Y-12729 | – | Unknown (received by NRRL from Guillermo Etienne Berumen, Instituto Mexicano del Petróleo) | 2 | 0 | 1 | 0 | 3 | 0.00 |
| Pp2 | NRRL Y-17741 | – | Emory oak (*Quercus emoryi*), Arizona, USA (W.T. Starmer) | 12,330 | 11,279 | 11,295 | 8,804 | 43,708 | 4.66 |
| Pp3 | NRRL Y-7556 (type strain) | CBS 2612 (type strain) | Black oak (*Quercus kelloggii*), California, USA (H.J. Phaff) | 2 | 0 | 2 | 0 | 4 | 0.00 |
| Pp4 | NRRL YB-378 | – | Elm (*Ulmus americana*), USA (L.J. Wickerham) | 11,735 | 10,752 | 10,862 | 8,489 | 41,838 | 4.46 |
| Pp5 | NRRL YB-4289 | – | Black oak (*Quercus kelloggii*), California, USA (H.J. Phaff) | 4,188 | 4,250 | 3,486 | 3,675 | 15,599 | 1.66 |
| GS115 | Derived from NRRL Y-11430 by mutagenesis (*his4*) | – | Commercial strain (Thermo Fisher Scientific) | 21 | 12 | 22 | 14 | 69 | 0.01 |
| CBS7435 | NRRL Y-11430 | CBS 7435 | Genome reference strain. US patent granted to Phillips Petroleum. | n/a | n/a | n/a | n/a | n/a | n/a |

NRRL, Northern Regional Research Laboratory, US Department of Agriculture. CBS, Centraalbureau voor Schimmelcultures (Westerdijk
Institute), The Netherlands.

**Fig 2.**
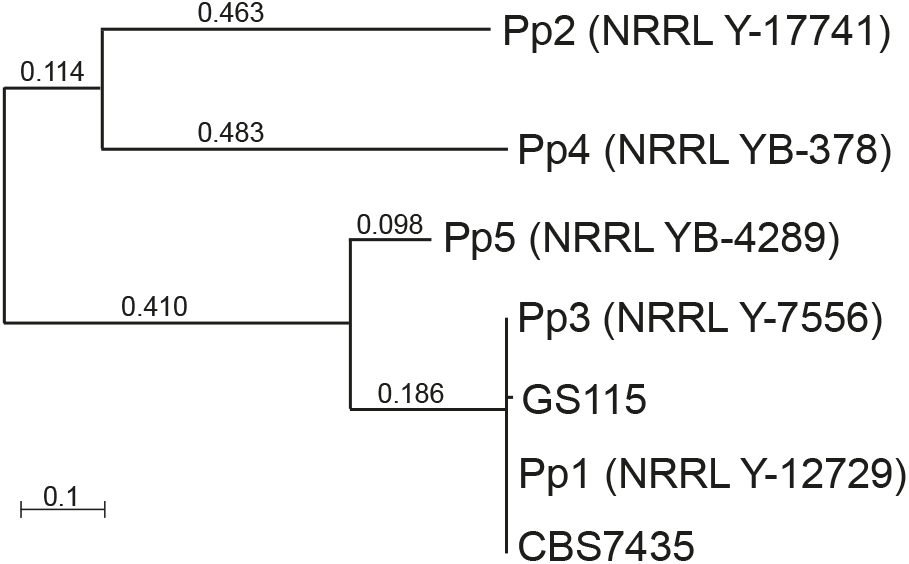
Phylogenetic tree of *K. phaffii* strains Pp1-Pp5, CBS7435 and GS115, based on 64,019 SNP sites. All non-trivial length branches have bootstrap values of 100%. Numbers on branches indicate branch lengths.

Strains Pp1 and Pp3 were found to be essentially identical to CBS7435, differing from it by only 3 or 4 nucleotides in the 9.4 Mb nuclear genome. Pp3 (NRRL Y-7556 / CBS2612) is a natural isolate from exudate of a black oak tree, and was designated by Kurtzman [32] as the type strain of *K. phaffii*. Pp1 and CBS7435, which were both deposited in culture collections by petroleum researchers (Table 1), seem to be duplicate accessions of this natural isolate. The mutagenized strain GS115 differs from its parent CBS7435 at 69 sites [5].

Among the three other natural isolates, Pp2 and Pp4 both show approximately 42,000– 44,000 SNP differences from the CBS7435 reference, and Pp5 shows approximately 16,000 (Table 1). Pp2 and Pp4 are also quite divergent from each other (Fig 2). The density of SNPs, at 1.66 – 4.66 SNPs per kb, is lower than the density seen in wild isolates of *S. cerevisiae* (e.g., the average nucleotide diversity among wild isolates from China is 8.08 differences/kb; [34]).

### Recovery of haploid segregants

We selected two of the most divergent strains from the phylogenetic tree, namely GS115 and Pp4, and crossed them in order to examine the meiotic recombination landscape in *K. phaffii*. GS115 has a *his4* mutation making it auxotrophic for histidine. We made a derivative of Pp4 that is auxotrophic for arginine (Pp4Arg^−^) by replacing the native *ARG4* gene with a *ble* cassette conferring resistance to zeocin (Zeo^R^). The *HIS4* and *ARG4* genes are both located on *K. phaffii* chromosome 1. They are 421 kb apart and on opposite sides of the centromere [33].

Diploid cells generated from the GS115 x Pp4Arg^−^ cross were identified by selecting for growth on media lacking His and Arg. Good mating was observed after incubating the cross for 3 days on diploid selection medium, resulting in a confluent patch of cells at the junction of streaked parental strains (**S1 Fig**). The diploid strain was sporulated and tetrad formation was confirmed by microscopy, but we were unable to dissect the tetrads using a Singer Sporeplay dissection microscope.

Because *K. phaffii* cells do not mate on rich (YPD) media, we reasoned that if an ascus is placed intact on YPD, so that its four spores germinate *in situ*, the resulting colony should contain a mixture of four different types of haploid cell that are the mitotic descendants of the four spores **(Fig 3)**. The four types of cell can be isolated simply by streaking out the original ascus-derived (mixed) colony, so that single cells initiate new colonies, each of which will have a homogeneous genotype that can be identified by replica plating onto appropriate media. Because our diploid was a double heterozygote (*HIS4/his4 ARG4/arg4*), it should produce three types of asci depending on how these markers segregate: parental ditypes (PD), non-parental ditypes (NPD), and tetratypes. In tetratype asci, which are formed if a single crossover occurs between the *HIS4* and *ARG4* loci, each spore has a different genotype and all four possible combinations of the two markers are present. Therefore, a tetratype ascus should produce four phenotypes after colonies are streaked out (His^+^ Arg^+^, His^−^ Arg^+^, His^+^ Arg^−^, and His^−^ Arg^−^), so we can identify colonies that are mitotic descendants of each of the four spores by replica plating onto appropriate media (Fig 3). In contrast, PD and NPD asci both produce only two colony phenotypes, so they cannot be used to identify descendants of all four spores.

**Fig 3.**
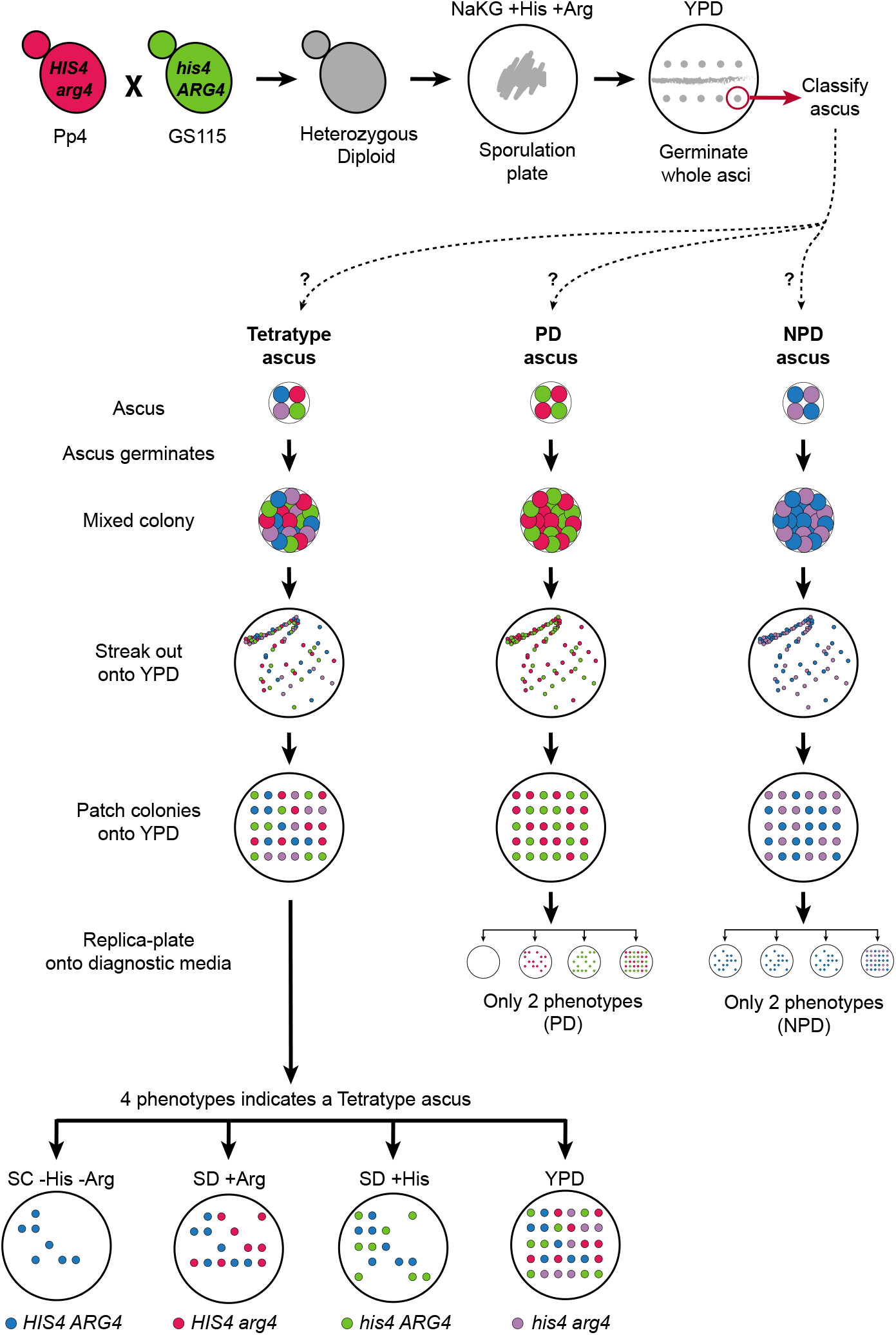
Experimental approach used to recover the four products of meiosis from an ascus without tetrad dissection. Parental strains Pp4 and GS115 were crossed to generate the resultant prototrophic heterozygous diploid. This diploid strain was sporulated under nitrogen-limiting conditions (NaKG +His +Arg medium). Whole asci containing four spores were picked up using a dissection microscope and placed on a YPD plate to germinate. The lower part of the diagram summarizes how asci were subsequently classified as tetratype, PD (parental ditype), or NPD (non-parental ditype) after streaking out and replica-plating. After germination, an ascus produces a mixed colony that contains a mixture of either 4 types of haploid cell (if the ascus is tetratype), or 2 types of haploid cell (if the ascus is PD or NPD), with respect to the *HIS4* and *ARG4* markers. After streaking out this mixed colony onto a fresh YPD plate and incubating for 2 days, single cells give rise to new colonies that each have a homogeneous genotype. These colonies are then patched in a grid pattern on another YPD plate, which is then replica-plated onto diagnostic media that allow the genotype of each colony to be inferred, hence enabling us to infer whether the ascus was tetratype, PD, or NPD. For asci inferred to be tetratype, the 4 types of haploid colonies (segregants) were retained for genome sequencing. For asci inferred to be PD or NPD, the segregants were discarded because it is not possible to identify all 4 products of the meiosis from their colony phenotypes. SC -His -Arg is synthetic complete media made without histidine and arginine. SD +Arg and SD +His are synthetic defined media supplemented with arginine or histidine. YPD is yeast peptone dextrose media (contains all amino acids).

Following this logic, we isolated four-spored asci from the sporulated culture using the micromanipulator, and placed them, without dissection, onto YPD agar so that each ascus germinated into a colony. We then streaked out these colonies to obtain new colonies initiated by single cells, patched the new colonies onto fresh YPD, and replica-plated them to assess their His and Arg phenotypes, looking for tetratypes. In this manner, we successfully recovered the four meiotic products from five tetratype asci (S1 Fig). Six other asci yielded three of the four expected phenotypes (‘trio’ asci), but we were unable to recover the fourth phenotype even after screening approximately 70 colonies from each of these asci. The absence of the last genotype in the trios is possibly due to epistatic interactions between loci from the Pp4 and GS115 genetic backgrounds, in the particular combinations that were formed in some spores, resulting in the failure of one spore to germinate.

### High-resolution mapping of meiotic recombination in *K. phaffii*

We sequenced the genomes of 38 segregants: 4 segregants from each of 5 tetrads, and 3 segregants from each of 6 trios. Each genome was sequenced to approximately 100x Illumina coverage, and the reads were used to genotype each segregant at every SNP site between the Pp4 and CBS7435 reference genome (which is almost identical to GS115; see Methods). Data analysis was carried out on 43,708 SNP markers. The median distance between consecutive markers in our cross is 96 bp, which is comparable to the *S. cerevisiae* cross analyzed by Mancera *et al*. (∼52,000 SNPs; median distance 78 bp) and the *Lachancea kluyveri* cross analyzed by Brion *et al*. (∼57,000 markers used; median distance 196 bp) [23, 25].

**Fig 4** shows a graphical representation of the segregation of SNP alleles on the four chromosomes in one tetrad (Tetrad 1), which has a total of 15 crossover events. As expected, a single crossover occurred in the interval between the *ARG4* and *HIS4* loci, producing the four genotypes of the tetratype ascus. With this data we were able to determine the locations of crossovers, and to identify gene conversion tracts such as the region of 3:1 segregation shown in the inset in Fig 4. Genotype calls from all tetrads and trios are given in **S1 Data**, and plots of their segregation patterns similar to Fig 4 are provided in **S2 Data**.

**Fig 4.**
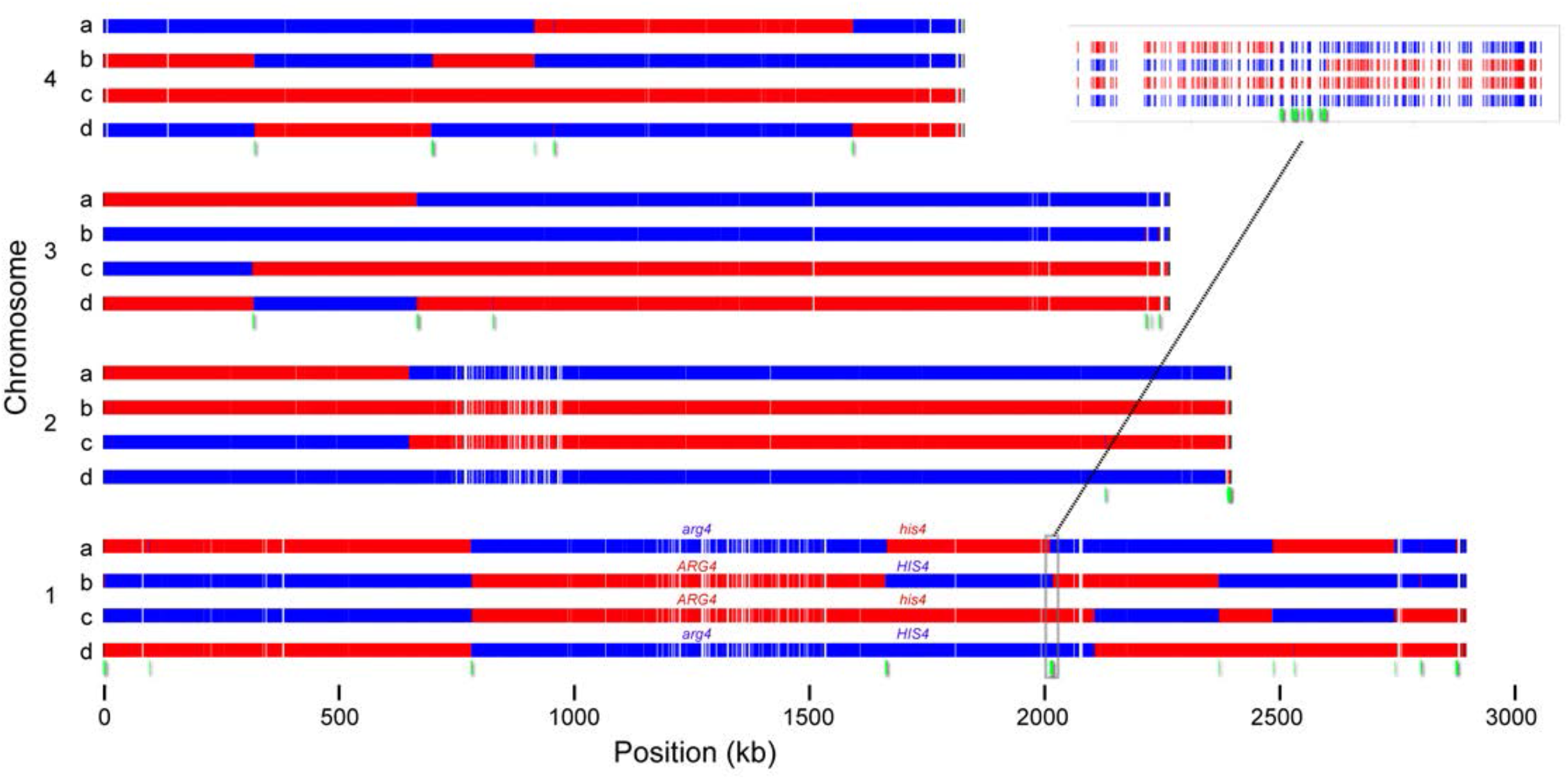
Segregation profile of one tetrad (Tetrad 1). Each group of four rows represents a single *K. phaffii* chromosome, indicated by numbers 1-4, and each row represents a spore, indicated by letters a-d. The four spores result from a single meiosis. Red segments represent regions inherited from the GS115 parent, and blue segments represent regions inherited from the Pp4 parent. *ARG4* and *HIS4* gene locations and genotypes are indicated. Red and blue bars are drawn at the location of every SNP difference between the two parental genomes. White gaps in the tracks are caused by stretches of DNA that are identical between the parents. Green segments below the chromosomes represent regions with non-Mendelian ratios (3:1 or 4:0) as illustrated in the inset.

We identified a total of 280 crossovers from the 11 meioses (**Table 2**; **S3 Data**). The mean number of crossovers per meiosis is 25.5, which is 3.5 times lower than the average in *S. cerevisiae* (90.5; [23]). The number varies about threefold (from 11 to 37) among the asci, with no systematic difference between tetrads and trios. The number of crossovers per meiosis in *K. phaffii* is also lower than in *S. paradoxus* (54.8), but similar to *Sch. pombe* (26.6) and *L. kluyveri* (19.9) [25, 27, 35].

**Table 2.** Summary of crossover (CO) counts in 11 *K. phaffii* asci.

| Ascus | Number of Crossovers |  |  |  |  |
| --- | --- | --- | --- | --- | --- |
|  | Chr. 1 | Chr. 2 | Chr. 3 | Chr. 4 | Whole Genome |
| *Tetrads* |  |  |  |  |  |
| Tetrad 1 | 8 | 1 | 2 | 4 | 15 |
| Tetrad 16 | 8 | 6 | 8 | 10 | 32 |
| Tetrad 20 | 7 | 4 | 4 | 8 | 23 |
| Tetrad 21 | 7 | 2 | 4 | 3 | 16 |
| Tetrad 27 | 7 | 6 | 5 | 6 | 24 |
| Total COs in 5 tetrads | 37 | 19 | 23 | 31 | 110 |
| COs per meiosis in 5 tetrads | 7.4 | 3.8 | 4.6 | 6.2 | 22.0 |
| *Trios* |  |  |  |  |  |
| Trio 10 | 10 | 5 | 5 | 12 | 32 |
| Trio 12 | 6 | 8 | 3 | 4 | 21 |
| Trio 19 | 12 | 12 | 11 | 0 | 35 |
| Trio 22 | 6 | 15 | 11 | 5 | 37 |
| Trio 31 | 1 | 2 | 7 | 1 | 11 |
| Trio 34 | 11 | 10 | 8 | 5 | 34 |
| Total COs in 6 trios | 46 | 52 | 45 | 27 | 170 |
| *All 11 asci* |  |  |  |  |  |
| Total COs in all 11 meioses | 83 | 71 | 68 | 58 | 280 |
| COs per meiosis in all 11 | 7.5 | 6.5 | 6.2 | 5.3 | 25.5 |
| Chromosome Length (kb) | 2895.354 | 2396.458 | 2263.458 | 1827.941 | 9383.211 |
| Crossovers/kb | 0.00261 | 0.00269 | 0.00273 | 0.00288 | 0.00271 |
| kb/crossover | 383.7 | 371.3 | 366.1 | 346.7 | 368.6 |

### Crossover frequency correlates with chromosome size

We found on average one crossover per 369 kb across the genome, a number that is consistent among the four chromosomes (Table 2). The average number of crossovers per meiosis on each chromosome in our data has a linear correlation with chromosome size (**Fig 5**), in agreement with the pattern seen in a variety of other fungal genomes [22, 25, 36, 37]. The trend line for *K. phaffii* has an intercept of 1.37 crossovers, in agreement with the occurrence one obligatory crossover per chromosome, which follows from the essential role that crossovers play in chromosome segregation. Every chromosome sustained at least one crossover in every meiosis (Table 2), except for chromosome 4 in Trio 19 for which all spores were derived from the Pp4 parent (i.e. 3:1 or 4:0 segregation) and thus did not have any crossovers.

**Fig 5.**
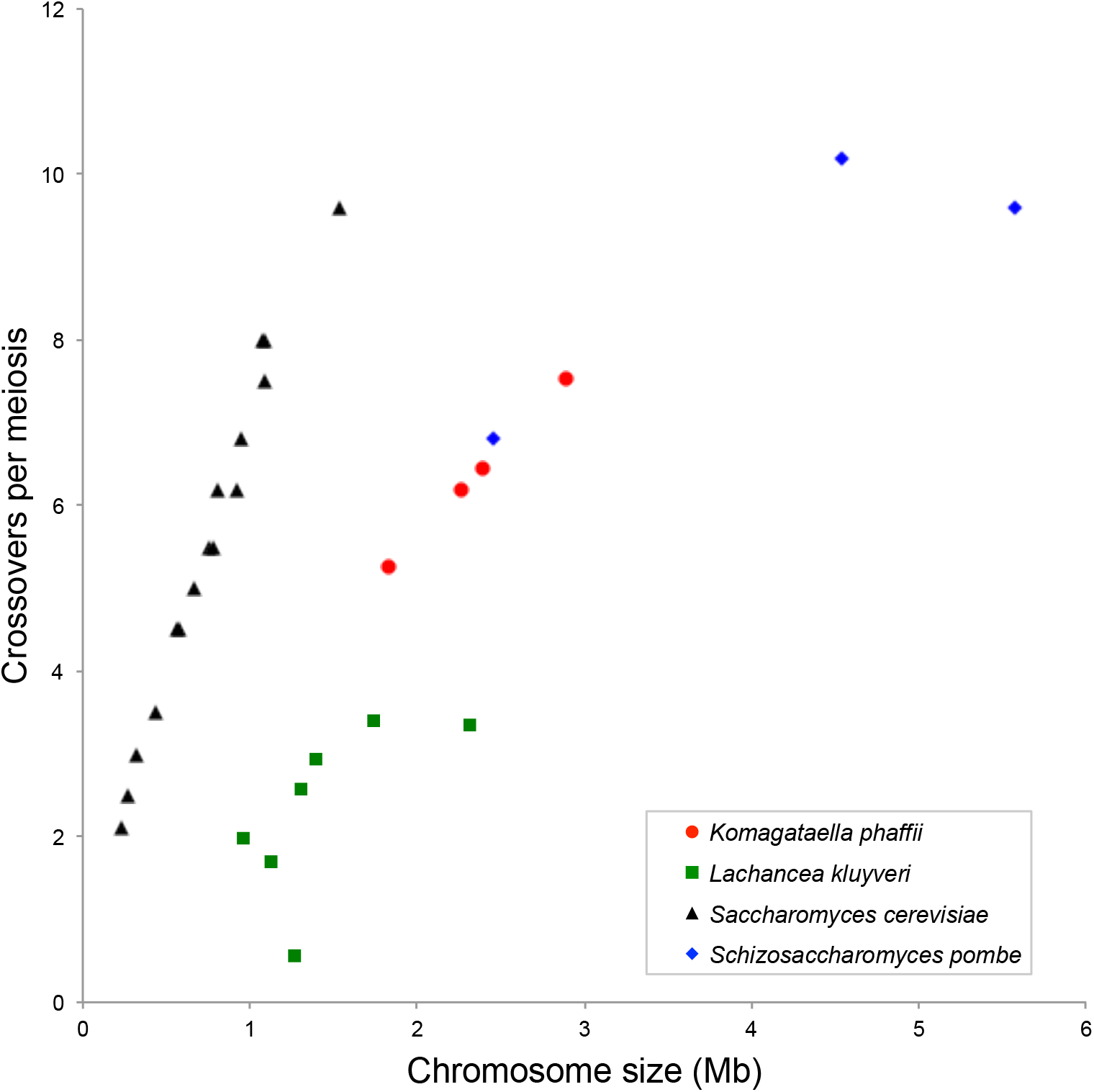
Relationship between chromosome size and the average number of crossovers per meiosis on that chromosome, in four ascomycete yeast species: *K. phaffii* (this study), *L. kluyveri* [25], *S. cerevisiae* [23], and *Sch. pombe* [35]. For *Sch. pombe*, we considered only the 10 F1 segregants in [35], omitting F2 segregants, and assumed that the actual number of crossovers in the ascus was twice the number of crossovers detected in the single segregant that was sequenced from each ascus; hence the results for *Sch. pombe* differ from [25]. The number of crossovers on the largest *Sch. pombe* chromosome may be abnormally low due to a large inversion [35]. For *L. kluyveri*, chromosome C has the lowest number of crossovers, and its low rate has been attributed to the origin of most of this chromosome from an ancient introgression event [25].

### Gene conversion tracts and non-crossover events

Gene conversion tracts are regions where 3:1 (or 4:0) segregation of alleles is seen. Gene conversion tracts that overlap with a crossover (‘crossover-associated’ tracts) are assumed to have arisen from heteroduplex DNA created at a double Holliday junction during formation of the crossover [38]. Gene conversion tracts that are not associated with crossovers are indicative of non-crossover (NCO) events, where an exchange between chromosomes was initiated but was resolved without causing a crossover. It should be noted that gene conversions and NCOs can only be detected if they extend over at least one site where there is a SNP difference between the parents, whereas crossovers can almost always be detected. It is also difficult to identify gene conversions and NCOs in data from trios, so we only examined the properties of NCOs in the five complete tetrads. The tetrads show an average of 17 NCOs and 22 crossovers per meiosis (Table 2; S3 Data). The ratio between crossovers and NCOs in *K. phaffii* is 1.3, which is lower than the previously reported ratios of 2.0 in both *S. cerevisiae* and *S. paradoxus* [23, 27], and 3.2 in *L. kluyveri* [25].

Examining the 110 crossovers in our five complete tetrads (Table 2) we observed 3:1 gene conversion tracts at 88 of them (80%; see examples for one tetrad in Fig 4), with only 22 crossovers not exhibiting tracts. For crossover-associated gene conversion tracts, an upper bound for the length of the tract can be obtained as the distance between the flanking SNPs that segregate 2:2. These upper bound lengths of crossover-associated gene conversion tracts range from 242 to 6412 bp, with a median of 1934 bp.

Gene conversion tracts are also frequently seen at the ends of chromosomes, and these subtelomeric tracts are generally longer than conversion tracts in the rest of the genome. The median lengths of gene conversion tracts are 1772 bp in subtelomeric regions vs. 695 bp in non-subtelomeric regions (*P* = 0.014 by two-tailed Wilcoxon rank-sum test), considering only the data from complete tetrads. The gene conversions in subtelomeric regions are likely the result of meiotic break-induced replication [23].

In *L. kluyveri*, Brion *et al*. [25] observed a high frequency of 4:0 segregation and ‘double crossovers’ (two crossovers at the same site, on two pairs of chromatids in a tetrad). They interpreted this pattern as as evidence of ‘return to mitotic growth’ (RTG), a process in which meiosis is initiated, abandoned, and then re-initiated after some rounds of mitotic division [25]. In contrast, we did not observe any evidence of double crossovers in our *K. phaffii* data, and found only one region with 4:0 segregation. There were two potential double crossover sites (Tetrad 1, chr. 1 at 781604-784124; and Tetrad 20, chr. 4 at 1125099-1131625), but after further examination we found each of them to be two independent single crossovers that were very close together. The Tetrad 1 site includes a region of 346 bp containing 9 SNP sites that segregate in a 4:0 pattern, probably due to the overlap of two adjacent 3:1 conversion tracts caused by two close but independent crossovers (Fig 4). We also observed four sites in which a crossover between two chromatids coincided with a patch of gene conversion on a third chromatid, a situation called ‘Type II tetrads’ by Liu *et al*. [27]. We scored these sites as both a crossover and an NCO (S3 Data).

### Recombination is suppressed in large regions around centromeres

Our experimental design made a crossover in the interval between the *ARG4* and *HIS4* markers on chromosome 1 obligatory in each of the 11 asci. In addition to these 11 obligatory crossovers, there was also a second crossover in this interval in two asci, involving a different pair of chromatids for each crossover. Looking at the locations of the 13 crossovers, we see that they are not randomly distributed along the 421-kb interval between *ARG4* and *HIS4*, which includes the centromere (*CEN1*). Instead, all the crossovers are close to *HIS4* and they seem to avoid the centromere region (**Fig 6A**). The hypothesis that crossovers are distributed uniformly in the interval is rejected by a statistical test (Kolmogorov-Smirnov test; *P* = 1.6e-6). The pattern suggests that crossing over near the centromere is suppressed.

**Fig 6.**
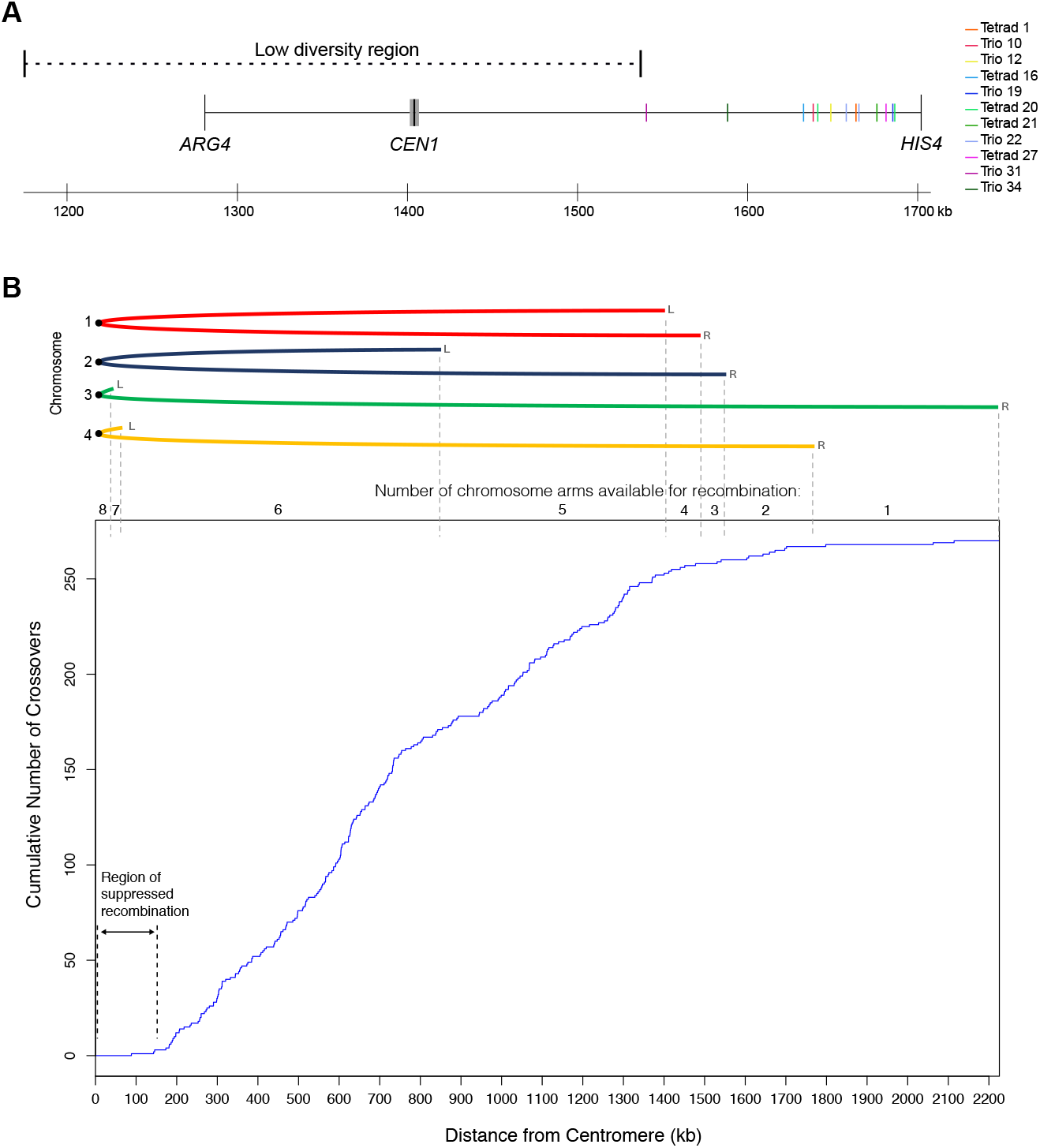
Suppressed recombination near *K. phaffii* centromeres. **(A)** Locations of crossovers in the *ARG4-HIS4* interval on chromosome 1. A crossover in this interval is obligatory in each of the 11 asci. Two asci (Tetrad 20 and Trio 22) had a second crossover in the interval, making 13 crossovers in total. The centromere (*CEN1*) is drawn to scale as a gray bar. The low diversity region around *CEN1* is indicated by a dashed line. **(B)** Distribution of distances from the centromere, for all 269 non-obligatory crossovers in our dataset. The graph shows the cumulative number of crossovers, plotted as a function of their distance from the centromere of the chromosome on which they occur. Data from all four chromosomes was pooled. The cartoon at the top shows the four chromosomes, folded at their centromeres, to show how many chromosome arms exist at any particular distance and hence are available for recombination (two of the eight chromosome arms are very short). Individual plots for each chromosome are shown in S2 Fig. Obligatory crossovers in the *ARG4-HIS4* interval were excluded from the analysis in B. For Tetrad 20 and Trio 22, we arbitrarily designated the more centromere-proximal crossover as non-obligatory and the more distal one as obligatory.

To investigate whether other crossovers are also suppressed near centromeres, we plotted the locations of all 269 non-obligatory crossovers in our dataset as a function of their distance from centromeres, both for the whole genome (**Fig 6B**), and for each chromosome individually (**S2 Fig**). The plots show that the recombination rate is low near centromeres. The distribution has an inflection point at a distance of approximately 150-200 kb from the centromere (Fig 6B), with only 3 crossovers occurring <150 kb from the centromere (the closest one is 88 kb away). Suppressed recombination around the centromere is seen on all four chromosomes (S2 Fig). *K. phaffii* has two metacentric chromosomes (chrs. 1 and 2), and two acrocentric chromosomes (chrs. 3 and 4) [39]. For the two metacentric chromosomes, suppression on both arms creates a zone of approximately 300 kb with low recombination.

### Low diversity in large regions around *CEN1* and *CEN2*

We found that the regions of suppressed recombination around the centromeres of chromosomes 1 and 2 coincide with regions of low sequence diversity among *K. phaffii* isolates. The low diversity regions are visible in Fig 4 as areas with many white gaps in the red and blue tracks, near the centers of chromosomes 1 and 2. The white gaps indicate places where there are no SNP differences between the parental strains GS115 and Pp4, i.e. they have identical sequences.

To examine this pattern in more detail, we plotted SNP diversity in the three natural isolates Pp4, Pp2 and Pp5, in 1-kb bins, relative to the CBS7435 reference genome (**Fig 7**). Pp4 and Pp2 both show large regions of low diversity around *CEN1* and *CEN2*, but not around the other two centromeres. The pattern in Pp5 is consistent, though this strain also shows several other large regions with high similarity to CBS7435. We defined low-diversity regions around *CEN1* and *CEN2* of Pp4 as the regions in which the average number of SNPs was less than 1 per kilobase – compared to a whole genome average of 4.5 per kilobase. The lengths of the low-diversity regions are 362 kb around *CEN1* and 240 kb around *CEN2*. This low diversity at centromeres in *K. phaffii* contrasts with the high variation seen in centromeric regions in fission yeast *Sch. pombe* [40].

**Fig 7.**
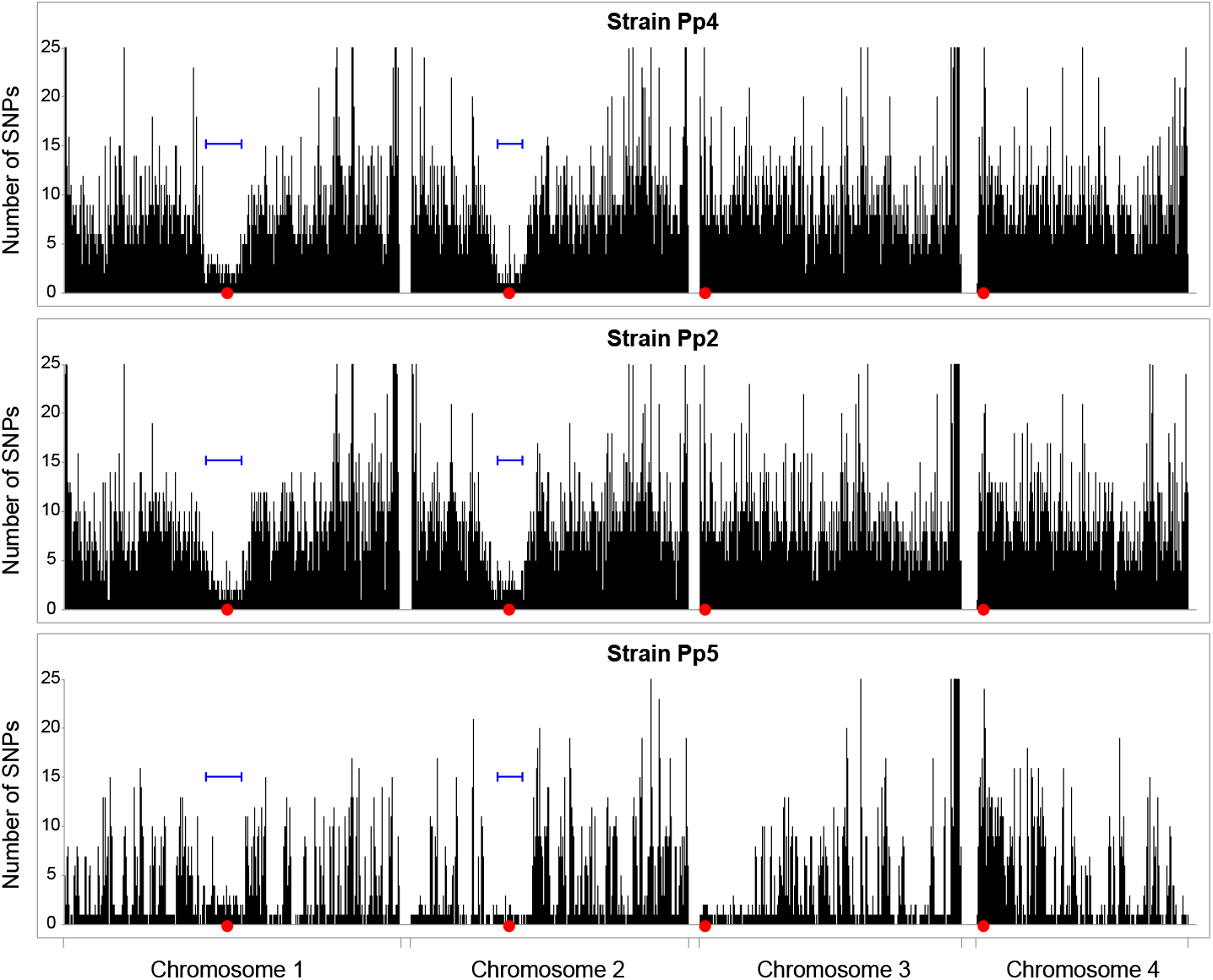
SNP density variation in natural isolates. For strains Pp4, Pp2, and Pp5, the number of SNP differences from CBS7435 in each 1-kb interval along each chromosome is plotted. Red dots mark centromeres, and blue bars mark the locations of the low-diversity regions that were defined in Pp4. The Y-axis maximum has been truncated to 25 SNPs/kb.

The low-diversity regions around *CEN1* and *CEN2* coincide approximately with their regions of suppressed recombination, but they are more than 20 times larger than the centromeres themselves. *K. phaffii* centromeres have a simple inverted repeat (IR) structure, with a total size of approximately 6 kb, and are located in non-transcribed regions of 6–9 kb that are flanked by protein-coding genes [39]. In contrast, the low-diversity regions in Pp4 and Pp2 extend outwards from the centromere for more than 100 kb in each direction (230 kb to the left and 132 kb to the right at *CEN1*; 110 kb to the left and 130 kb to the right at *CEN2*). There are many protein-coding genes in the low-diversity regions, including *ARG4*, and the gene density is the same as in the rest of the genome. All the crossovers in the *ARG4-HIS4* interval occurred outside the low-diversity region around *CEN1* (Fig 6A).

## Discussion

We found that *K. phaffii* has a recombination rate approximately 3.5 times lower than in *S. cerevisiae*. This result is not very surprising, because *S. cerevisiae* is one of the most recombinogenic species known. Compared to other yeasts, the *S. cerevisiae* genome is organized into chromosomes that are shorter and more numerous, as a result of its whole-genome duplication. When chromosome size is taken into account, the numbers of crossovers per megabase per meiosis in *K. phaffii* are similar to the numbers in *L. kluyveri* and *Sch. pombe* (i.e. the points are on the same line in Fig 5), but substantially lower than in *S. cerevisiae*.

The most surprising result from our analysis was the discovery of a very large region of suppression of meiotic recombination around all four *K. phaffii* centromeres. Recombination is also suppressed close to centromeres in *S. cerevisiae*, but the effect only extends over approximately 10 kb [23, 37, 41], compared to approximately 300 kb in *K. phaffii*. In *S. cerevisiae*, crossovers near centromeres are suppressed by the kinetochore, which attaches to a single nucleosome that contains centromeric histone H3 (Cse4), and by cohesin bound to the pericentromeric region [41–43]. At *K. phaffii* centromeres, the region of Cse4 binding is less than 10 kb long [39], so the reason why recombination is suppressed over a 300 kb region is unclear, but suggestive of a possible difference in the extent of pericentric cohesin loading between the point centromeres of *S. cerevisiae* and the IR-containing centromeres of *K. phaffii*.

On the two metacentric chromosomes (chrs. 1 and 2), the region of low recombination around the centromere coincides with a region of low sequence diversity. The pattern on these chromosomes is consistent with previous observations of low sequence diversity in non-recombining regions of animal [44, 45] and plant [46] genomes, and has been attributed to background selection. In background selection, when deleterious mutations arising in regions of low recombination are eliminated by natural selection, any linked neutral variants are also eliminated and hence genomic diversity in the region is reduced [47]. In general, genomic diversity is positively correlated with the recombination rate among regions of eukaryotic genomes, though this pattern is only weakly seen in *S. cerevisiae* [48, 49]. In contrast to the metacentric chromosomes, the centromere regions of the acrocentric *K. phaffii* chromosomes (chrs. 3 and 4) show low recombination without low sequence diversity. Chromosome 4 may be a special case, because *CEN4* is located approximately in the center of a 138-kb region that is inverted between cells of mating types **a** and α, so the region is in opposite orientations on the two copies of chromosome 4 in diploid cells. A single crossover near *CEN4* would generate inviable chromosomes with huge deletions or duplications [12, 39], and we did not see any crossovers in the 138-kb region. Nevertheless, nucleotide diversity in this region is high.

Our work provides a foundation for future quantitative trait locus analysis in *K. phaffii*. Because most *K. phaffii* strains used in industry are derivatives of a single isolate, CBS7435, it may be possible to identify beneficial alleles in other isolates of this species and introduce them into commercial strains, either by crossing or by genome editing. However, we also found that the pool of potential alleles currently available is quite small, because only a few natural isolates of *K. phaffii* are known.

Our method of tetrad analysis without tetrad dissection enabled us to study the genomic recombination landscape at high resolution in a species for which the only previous method of genetic analysis was random spore analysis. Tetrad analysis is valuable as a method for studying aspects of the recombination process that are not otherwise accessible, such as a genome-wide view of gene conversion events [21]. Our experiments forced a crossover to occur in one genomic interval, but this was not an essential part of the design and we could have used selectable markers on two different chromosomes. Our method for recovering segregants from tetratype asci is applicable to any organism in which the unseparated segregants from a meiosis can be recovered as a unit, provided that the segregants do not mate spontaneously with each other.

## Materials and methods

### Strain generation and growth conditions

The NRRL strains of *K. phaffii* used in this study (Table 1) were obtained from the USDA NRRL Culture Collection (USA). Strain CBS7435 was obtained from the Spanish Type Culture Collection as CECT 11047. All strains were grown in YPD (1% yeast extract, 2% peptone, 2% glucose) medium at 30 °C and 200 rpm of agitation.

Parental strains used to generate the recombination map were the laboratory strain GS115 (Thermo Fisher Scientific) and the natural isolate Pp4 (NRRL YB-378). GS115 is auxotrophic for histidine (His^−^) due to a C557R point mutation in *HIS4* [5, 8, 50], and we confirmed this by resequencing the GS115 genome. Pp4 was made auxotrophic for arginine (Arg^−^) by replacing *ARG4* [51] with the drug marker *ble* conferring zeocin resistance. The pILV5-*ble* cassette (ZEO) from *K. phaffii* plasmid pJ902-15 (Atum, USA) was flanked by 1 kb of right- (HR) and 1 kb of left- (HL) homology arms originating from the sequences flanking the *ARG4* gene of Pp4 (**S4 Data**). These three fragments were PCR-amplified individually (New England Biolabs Q5 high-fidelity 2X master mix, 56 °C annealing temperature, 25 cycles) and joined in a single step by overlap extension PCR (1^st^ step: equimolar mix of PCR fragments ZEO, HR and HL, NEB Q5 high-fidelity 2X master mix, 60 °C annealing temperature, 12 cycles, no primers added; 2^nd^ step: addition of primers for fragment HL+ZEO+HR, 67 °C annealing temperature, 25 cycles) to produce a 3.5 kb insert flanked by *Sac*I restriction sites. The HL+ZEO+HR insert was ligated into the unique *Sac*I site in plasmid pUC57 (Thermo Fisher Scientific) and transformed into One Shot TOP10 Chemically Competent *E. coli* cells (Thermo Fisher Scientific) generating the 6.25 kb ARG4del plasmid (S4 Data). This plasmid was digested with the restriction enzyme pair *Bse*RI-*Afl*II for homologous recombination into Pp4, replacing the native *ARG4* gene with the ZEO construct. Transformation was carried out by electroporation following Atum’s guidelines, and transformant colonies harboring the *ARG4* deletion (*ARG4*Δ::*ble*) were selected on YPD plates supplemented with zeocin (200 μg ml^−1^). The Pp4Arg^−^ mutant phenotype was confirmed by absence of growth in Synthetic Defined (SD) minimal medium (2% glucose, 6.7 g l^−1^ yeast nitrogen base without amino acids) and by PCR amplification of the targeted locus.

The cross of parental strains GS115 and Pp4Arg^−^ was carried out by making parallel streaks of the two parental strains on YPD, and then velvet replica plating these streaks onto a mating plate twice at right angles so that they intersected as a grid [10]. The mating plate contained NaKG agar media (0.5% sodium acetate, 1% potassium chloride, 1% glucose, 2% bacto agar) plates supplemented with L-histidine (50 mg l^−1^) and L-arginine (100 mg l^−1^) (NaKG +His +Arg). The mating plate was incubated for 2 days at room temperature and subsequently replica plated onto diploid selection medium (SC -His -Arg: SD medium supplemented with -Arg/-His drop-out mix). Diploids were incubated for 3 days at 30 °C, and streaked for a second time for phenotype confirmation and generation of single colonies.

### Recovery of asci and haploid segregants

Sporulation was induced by streaking diploid cells on NaKG+His+Arg plates and incubating for ≥5 days at room temperature. A loop of sporulated material was resuspended in sterile distilled water and digested for 10 min at 30 °C with Zymolyase 100T (Stratech). The mixture was washed and resuspended in sterile distilled water, and a 10 μl drop of this suspension was left to run across a YPD plate, forming a horizontal streak. The plate was incubated for 4 hours at 30 °C to allow asci to swell and become more visible under the 40X lens of a SporePlay+ dissection microscope (Singer Instruments). Whole asci were picked up and placed individually on the YPD plate, which was incubated for 2 days at 30 °C until colonies were visible. Each colony starting from a single ascus was picked and streaked onto a fresh YPD plate for single colonies (segregants). Approximately 20 to 70 segregants per ascus were picked and patched on a fresh YPD plate, incubated at 30 °C overnight, and replica plated onto a set of four diagnostic plates for identification of each segregant type (Fig 3; S1 Fig). The diagnostic plates contained: SC -His -Arg media (only *HIS4 ARG4* segregants grow); SD +His media (*his4 ARG4* and *HIS4 ARG4* segregants grow); SD +Arg media (*HIS4 arg4* and *HIS4 ARG4* segregants grow); and YPD (*his4 arg4* and all other segregants grow). By visual inspection and comparison of the colonies that grew on each media it was possible to identify the 4 (or 3) meiotic products originating from tetratype asci. Asci that yielded only two phenotypes were considered to be PD or NPD and discarded. It is important to note that *K. phaffii* spores tend to clump together, which can be noticed by the high number of colonies showing prototrophic phenotypes (S1 Fig; [10]). Therefore segregants from all candidate tetratype asci were re-patched onto all diagnostic media to confirm their phenotypes before genome sequencing.

### Genomic DNA extraction and genome sequencing

Genomic DNA was extracted from stationary-phase cultures by homogenization with glass beads followed by phenol-chloroform extraction and ethanol precipitation. Purified DNA was concentrated with the Genomic DNA Clean & Concentrator-10 (Zymo Research). Natural isolates Pp1-Pp5 were sequenced at the University of Missouri core facility using an Illumina HiSeq 2500 instrument, with single-end 51 bp reads, to approximately 30x coverage, and assembled using SPAdes version 3.10 [52]. The parental strains used in the cross (Pp4Arg^−^ and GS115) and the 38 segregants, were each sequenced to approximately 100x coverage (paired-end 150 bp reads) on an Illumina HiSeq 4000 by BGI Tech Solutions (Hong Kong).

### SNP calling

BAM alignments of Illumina reads from each segregant to the CBS7435 reference genome [33] were generated using the Burrows-Wheeler Aligner (BWA) with default parameters [53]. Unmapped reads were removed using SAMtools [54] and headers were added using the AddOrReplaceReadGroups program in Picard Tools [http://picard.sourceforge.net]. Variants against the reference were called with the GATK HaplotypeCaller tool in DISCOVERY genotyping mode with the following parameters: “-stand_emit_conf 10 -stand_call_conf 30 -- min_base_quality_score 20 emitRefConfidence GVCF”. For the segregants from each tetrad and trio these GVCF files were then combined using GATK into joint genotype calls for each tetrad/trio. For improved clarity in downstream analysis the 69 SNPs between CBS7435 and GS115 [5] were removed from these lists – they are an artefact of our decision to use CBS7435 as the reference genome for alignment (because it had better assembly and annotation), whereas GS115 was the actual parent in our cross.

### Crossover detection

A bespoke Java program was used, taking the joint genotype calls as input, to list the genotype of the 4 segregants in each tetrad (or 3 segregants in each trio) at every SNP site (S1 Data). Genotypes of segregants at SNP sites were coded as 0 (for a nucleotide matching GS115) or 1 (for a nucleotide matching Pp4). Sites called as indels or heterozygous were discarded. Genotype lists for each tetrad were input into the plotting tool in the ReCombine package [55] to make red/blue maps of the segregation patterns in asci (Fig 4; S2 Data), after modifying the tool to plot 4 chromosomes instead of 16. Crossover and noncrossover events were identified using a program to find every SNP site in a trio/tetrad whose segregation pattern was different from neighboring site(s), and scoring manually. We ignored noncrossovers involving only a single SNP site, and ignored the subtelomeric regions of each chromosome (Chr. 1: coordinates <8 kb and >2861 kb; Chr. 2: <12 kb and >2373 kb; Chr. 3: >2168 kb; Chr. 4: <2 kb and >1795 kb) except where stated. For analysis of recombination rates near centromeres, we calculated distances from centromeres as the distance from the midpoint of the unique central region between the IRs: coordinates 1404539 (*CEN1*), 847809 (*CEN2*), 37576 (*CEN3*), and 61908 (*CEN4*) in the reference genome sequence of Sturmberger *et al*. [33].

### Phylogenetic tree

The phylogenetic tree of strains CBS7435, GS115 and Pp1-Pp5 was generated using PhyML (HKY model) [56] from the 64,019 SNP sites that vary among these strains, calculated using GATK (as described above for the segregants) for Pp1-Pp5 compared to CBS7435, and using MUMmer’s *nucmer* whole genome alignment program [57] for the GS115 to CBS7435 comparison.

## Acknowledgments

We thank Darren Nesbeth and Ben Offei for advice, the NRRL and CECT culture collections for strains, Geraldine Butler and Wolfe lab members for comments on the manuscript, and Nadia Trigoub-Browne for translation.

## Author contributions

**Conceptualization:** Stephanie Braun-Galleani, Kenneth H. Wolfe

**Formal analysis:** Julie A. Dias, Adam P. Ryan, Kevin P. Byrne, Kenneth H. Wolfe

**Funding acquisition:** Kenneth H. Wolfe

**Investigation:** Stephanie Braun-Galleani, Julie A. Dias, Aisling Y. Coughlan

**Methodology:** Stephanie Braun-Galleani, Julie A. Dias

**Resources:** Kevin P. Byrne, Kenneth H. Wolfe

**Supervision:** Kevin P. Byrne, Kenneth H. Wolfe

**Writing – original draft:** Stephanie Braun-Galleani, Julie A. Dias

**Writing – review & editing:** Kevin P. Byrne, Kenneth H. Wolfe

## Supporting information

**S1 Fig. Crossing of parental strains and identification of segregant genotypes.** (A) Crossing of parental strains. Parallel streaks of the two parental strains Pp4Arg^−^ and GS115 were made on a YPD plate, and then replica-plated twice perpendicularly onto a mating plate (NaKG +His +Arg) to make a grid pattern. After growth, the mating plate was then replica plated onto the SC -His -Arg selection plate shown. Diploid cells (His^+^ Arg^+^) grow at the intersections of the streaks. (B) Initial identification of segregants from Tetrad 27, by replica plating onto 4 media. An example of each class of segregant is circled. (C) Verification of segregant genotypes from Tetrad 27 by a second round of replica plating. (PDF)

**S2 Fig. Locations of crossovers on each chromosome, as a function of their distance from the centromere.** Red lines and numbers 1-2 indicate the number of chromosome arms that exist at any particular distance. All chromosomes are plotted on the same horizontal scale. Obligatory crossovers in the *ARG4-HIS4* interval on chromosome 1 were excluded from this analysis (for Tetrad 22 and Trio 20, which have two crossovers in this interval, we arbitrarily designated the more centromere-proximal crossover as non-obligatory and the more distal one as obligatory). (PDF)

**S1 Data. Genotype calls at SNP sites in each ascus.** Values of “0” indicate the same nucleotide as in GS115, and “1” indicates a match to Pp4. Spores were designated as types **a** (*his4 arg4*), **b** (*HIS4 ARG4*), **c** (*his4 ARG4*) or **d** (*HIS4 arg4*). All four types of spore were recovered from tetrads; Trios 10, 22 and 19 lack a spore of type **a**, and Trios 31, 34 and 12 lack a spore of type **d**. Chromosome coordinates are from the CBS7435 reference genome assembly of Sturmberger *et al*. [33], NCBI accession numbers LT962476.1 to LT962479.1. (ZIP)

**S2 Data. Segregation profiles of all *K. phaffii* tetrads and trios analyzed, as in Fig 4.** (ZIP)

**S3 Data.** Details of crossover and noncrossover sites identified in each tetrad and trio. (XLS)

**S4 Data.** Nucleotide sequence of the ARG4del plasmid used to delete *ARG4* in strain Pp4 (TXT)

